# *ReCSAI*: Recursive compressed sensing artificial intelligence for confocal lifetime localization microscopy

**DOI:** 10.1101/2022.05.06.490886

**Authors:** Sebastian Reinhard, Dominic A. Helmerich, Dominik Boras, Markus Sauer, Philip Kollmannsberger

**Affiliations:** Department of Biotechnology and Biophysics, Biocenter, University of Wuerzburg, 97074 Wuerzburg, Germany; Center for Computational and Theoretical Biology, University of Wuerzburg, 97074 Wuerzburg, Germany

**Keywords:** Compressed sensing, AI, SMLM, FLIMbee

## Abstract

Localization-based super-resolution microscopy resolves macromolecular structures down to a few nanometers by computationally reconstructing fluorescent emitter coordinates from diffraction-limited spots. The most commonly used algorithms are based on fitting parametric models of the point spread function (PSF) to a measured photon distribution. These algorithms make assumptions about the symmetry of the PSF and thus, do not work well with irregular, non-linear PSFs that occur for example in confocal lifetime imaging, where a laser is scanned across the sample. An alternative method for reconstructing sparse emitter sets from noisy, diffraction-limited images is compressed sensing, but due to its high computational cost it has not yet been widely adopted. Deep neural network fitters have recently emerged as a new competitive method for localization microscopy. They can learn to fit arbitrary PSFs, but require extensive simulated training data and do not generalize well. A method to efficiently fit the irregular PSFs from confocal lifetime localization microscopy combining the advantages of deep learning and compressed sensing would greatly improve the acquisition speed and throughput of this method. Here we introduce *ReCSAI*, a compressed sensing neural network to reconstruct localizations for confocal *d*STORM, together with a simulation tool to generate training data. We implemented and compared different artificial network architectures, aiming to combine the advantages of compressed sensing and deep learning. We found that a U-Net with a recursive structure inspired by iterative compressed sensing showed the best results on realistic simulated datasets with noise, as well as on real experimentally measured confocal lifetime scanning data. Adding a trainable wavelet denoising layer as prior step further improved the reconstruction quality. Our deep learning approach can reach a similar reconstruction accuracy for confocal *d*STORM as frame binning with traditional fitting without requiring the acquisiton of multiple frames. In addition, our work offers generic insights on the reconstruction of sparse measurements from noisy experimental data by combining compressed sensing and deep learning. We provide the trained networks, the code for network training and inference as well as the simulation tool as python code and Jupyter notebooks for easy reproducibility.

## Introduction

The resolution of classical fluorescence microscopy is limited by the Abbe criterion (1). In the past decades, several super-resolution techniques to surpass this limit have been developed. One of them is single molecule localization microscopy (SMLM), which is based on localizing the position of individual fluorescent dyes. By fitting a model of the theoretical photon distribution, the point spread function (PSF), to the measured signal, the emitter position can be precisely determined (2). While this problem was quickly solved for perfect samples, reality is often more difficult. Overlapping or varying photon distributions as well as low signal-to-noise ratio still pose a challenge. Various approaches are used to reconstruct super-resolved positions of individual emitters, such as intensity centroids, fitting Gaussian or more complex (e.g. Zernike polynomial) functions (3), compressed sensing (4, 5), and deep neural networks (6, 7).

An interesting application of SMLM is the simultaneous imaging of different targets using multiple colors however, suitable fluorescent dyes are very limited, and chromatic aberrations are unavoidable for different emission wavelength (8). A promising workaround is to distinguish dyes with similar emission wavelength by their different lifetime (9). This detection method is based on confocal scanning, where a laser scans the sample with a sampling rate similar to the switching rate of fluorescent dyes. This introduces distorted and disrupted PSFs that cannot be properly localized by fitting a parametric PSF model. To solve this problem, Thiele et al.(9) acquired multiple frames, projected them onto each other to obtain complete PSFs, and applied conventional fitting. An efficient method to fit the irregular, chopped PSFs in individual frames would greatly improve the acquisition speed and throughput of confocal lifetime localization microscopy.

The nonlinearities of irregular, chopped PSFs require a large degree of flexibility in the fitting function while maintaining high precision. Most classical algorithms are based on fitting a Gaussian or similar function to the measured photon distribution for each individual emitter. While these methods approach the theoretical lower bound for individual emitters, they fail when emitters overlap or the PSF is irregular. For overlapping emitters at high density, compressed sensing (CS) is superior to conventional fitting (10). CS works by solving the inverse problem of recovering a super-resolved image of the emitters from a noisy, low-resolution measurement using sparsity in the spatial (4, 5) or correlation (11) domain as constraint. Due to its high computational demands, CS has so far not found widespread use. Artificial Neural Networks (ANN) are well suited for fitting complex PSFs, as they are essentially high-dimensional function approximators. Recently, ANN-based fitters such as DeepSTORM (6) or DECODE (7) achieved outstanding results in a SMLM reconstruction benchmark (10), beating CS at high emitter density. Since the iterative process of compressed sensing can be expressed as a differentiable operation, it should be possible to integrate it into a neural network to combine the advantages of both approaches.

Here, we present a novel trainable fitting algorithm that combines wavelet denoising, compressed sensing, and deep learning to recover emitter locations from confocal *d*STORM data with chopped, irregular PSFs. We developed a simulator producing accurate ground truth for training, including various distortions and noise as well as temporal context over multiple frames. We then trained different neural networks combining existing approaches with a novel trainable CS layer and wavelet filters, and evaluated and compared their performance on simulated and experimental data using common accuracy metrics as well as Fourier Ring Correlation (12) and LineProfiler (13).

## Methods

### Antibody labeling

For antibody labeling, an excess of Cy5-NHS (GE-Healthcare, PA15101) was used. Goat anti-rabbit IgG (Invitrogen, 31212) was used as secondary antibody for microtubules in FLIM experiments. Antibody labeling was performed at room temperature for 4h in labeling buffer (100 mM sodium tetraborate (Fulka, 71999), pH 9.5) following the manufacturers standard protocol. Briefly, 100 μg antibody were reconstituted in labeling buffer using 0.5 ml spin-desalting columns (40K MWCO, ThermoFisher, 87766). An 5x excess of Cy5-NHS (GE-Healthcare, PA15101) was used. Antibody conjugates were purified and washed up to three times using spin-desalting columns (40K MWCO) in PBS (Sigma-Aldrich, D8537-500ML) to remove excess dyes. Finally, antibody concentration and DOL were determined by UV-vis absorption spectrometry (Jasco V-650).

### Cell culture

African green monkey kidney fibroblast-like cells (COS7, Cell Lines Service GmbH, Eppelheim, #605470) were cultured in DMEM (Sigma, #D8062) containing 10 % FCS (Sigma-Aldrich, #F7524), 100 U/ml penicillin and 0.1 mg/ml streptomycin (Sigma-Aldrich, #P4333) at 37 °C and 5 % CO_2_. Cells were grown in standard T25-culture flasks (Greiner Bio-One).

### Immunostaining

For immunostaining, cells were seeded at a concentration of 2.5 × 10^4^ cells/well into 8 chambered cover glass systems with high performance cover glass (Cellvis, C8-1.5H-N) and stained after 3 hours of incubation at 37 °C and 5 % CO_2_. For microtubule immunostaining, cells were washed with pre-warmed (37 °C) PBS (Sigma-Aldrich, D8537-500ML) and permeabilized for 2 min with 0.3 % glutaraldehyde (GA) + 0.25 % Triton X-100 (EMS, 16220 and ThermoFisher, 28314) in pre-warmed (37° C) cytoskeleton buffer (CB), consisting of 10 mM MES ((Sigma-Aldrich, M8250), pH 6.1), 150 mM NaCl (Sigma-Aldrich, 55886), 5 mM EGTA (Sigma-Aldrich, 03777), 5 mM glucose (Sigma-Aldrich, G7021) and 5 mM MgCl_2_ (Sigma-Aldrich, M9272). After permeabilization, cells were fixed with a pre-warmed (37° C) solution of 2 % GA for 10 min. After fixation, cells were washed twice with PBS (Sigma-Aldrich, D8537-500ML) and reduced with 0.1 % sodium borohydride (Sigma-Aldrich, 71320) in PBS for 7 min. Cells were washed three times with PBS (Sigma-Aldrich, D8537-500ML) before blocking with 5 % BSA (Roth, #3737.3) for 30 min. Subsequently, microtubule samples were incubated with 2 ng/μl rabbit anti-*α*-tubulin primary antibody (Abcam, #ab18251) in blocking buffer for 1 hour. After primary antibody incubation, cells were rinsed with PBS (Sigma-Aldrich, D8537-500ML) and washed twice with 0.1 % Tween20 (ThermoFisher, 28320) in PBS (Sigma-Aldrich, D8537-500ML) for 5 min. After washing, cells were incubated in blocking buffer with 4 ng/μl of dye-labeled goat antirabbit IgG secondary antibodies (Invitrogen, 31212) for 45 min. After secondary antibody incubation, cells were rinsed with PBS (Sigma-Aldrich, D8537-500ML) and washed twice with 0.1 % Tween20 (ThermoFisher, 28320) in PBS (Sigma-Aldrich, D8537-500ML) for 5 min. After washing, cells were fixed with 4% formaldehyde (Sigma-Aldrich, F8775) for 10 min and washed three times in PBS (Sigma-Aldrich, D8537-500ML) prior to imaging.

### Fluorescence Lifetime Imaging Microscopy (FLIM)

All single molecule fluorescence lifetime measurements were performed on a MicroTime200 (PicoQuant, Berlin, Germany) time-resolved confocal fluorescence microscope setup (Figure 1 a) consisting of a FLIMbee galvo scanner (Pico-Quant, Berlin, Germany), an Olympus IX83 microscope including an oil-immersion objective (60×, NA 1.45; Olympus), 2 single photon avalanche photodiodes (SPAD) (Excelitas Technologies, 75154 K3, 75154 L6) and a TimeHarp300 dual channel board. Pulsed excitation was performed using a white-light laser (NKT photonics, superK extreme) which was coupled into the MicroTime200 system via a glass fiber (NKT photonics, SuperK FD PM, A502-010-110). A 100 μm pinhole was used for all measurements. The emission light was split onto the SPADs using a 50:50 beamsplitter (PicoQuant, Berlin, Germany). To filter out afterglow effects of the SPADs as well as scattered and reflected light, two identical bandpass filters (ET700/75 M, Semrock, 294808) were installed in front of the SPADs. The measurements were performed and analyzed with the SymPhoTime64 software (PicoQuant, Berlin, Germany). The microtubule measurements were performed with an irradiation intensity of 5 kW cm^−2^ in T3 mode with 25 ps time-resolution. The pixel dwell time was 100 μs and a monodirectional line frequency of 108.7 Hz was used. The corresponding frame frequency was 2.4 Hz. Measurements were performed in PBS-based photoswitching buffer containing 100 mM *β*-mercaptoethylamine (MEA, Sigma-Aldrich) adjusted to pH 7.6.

**Fig. 1.**
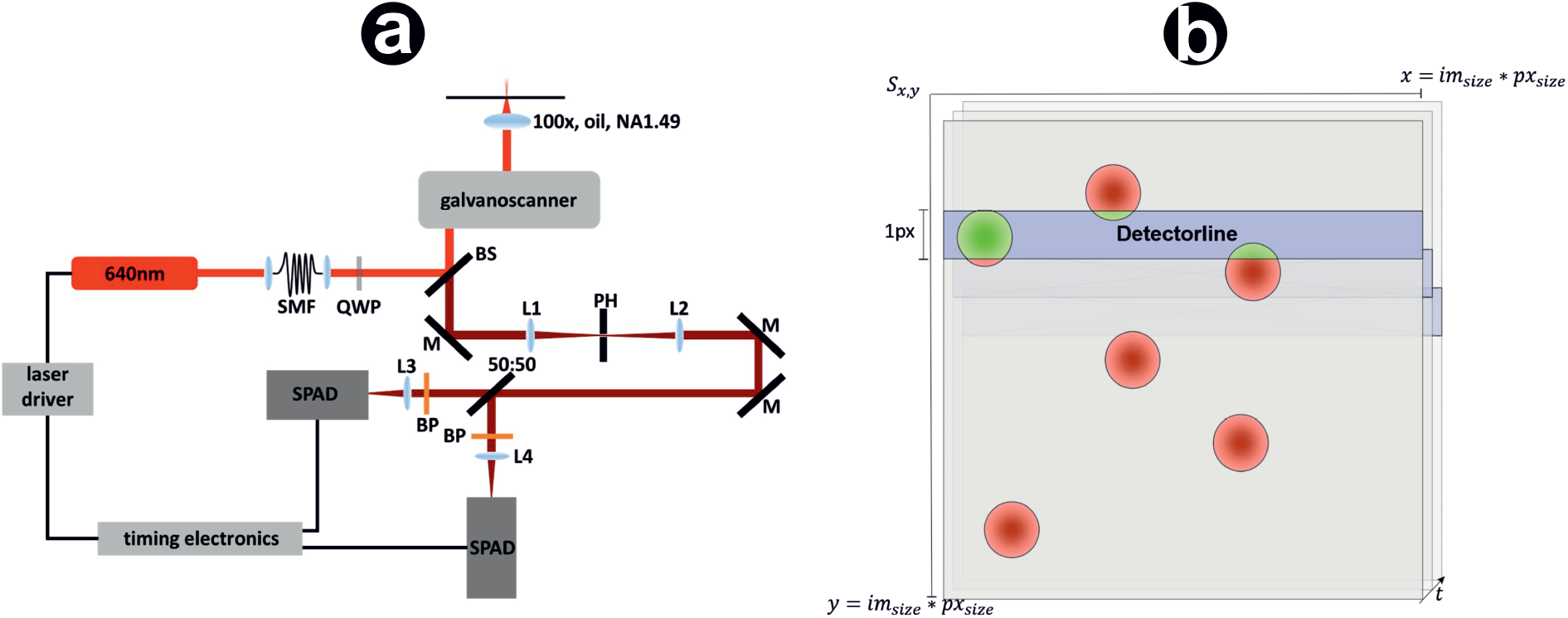
Confocal *d*STORM data acquisition process and data simulation. (a) The pulsed 640 nm excitation light is converted to radial polarisation with a quarter wave plate (QWP) after passing through a single mode fibre (SMF), reflected by a beam splitter (BS) into a galvanometric laser scanner and focused by an oil immersion objective. The collected fluorescent emission from the sample is descanned, passed through the BS, reflected by mirrors (M) and focused onto the pinhole (PH), then onto the single-photon avalanche photodiodes (SPAD) using lenses (L1, L2, L3 and L4). Band pass filters (BP) block scattered excitation light and prevent afterglow effects of the detectors. (b) The galvo scanner is a one-pixel scanner rastering the image line by line. At time *t*, only the blue marked part of *S* is active, representing the horizontal acquisition line of the FLIMbee detector. The acquisition speed in x-direction is sufficiently fast to be neglected during simulations. Only active fluorophores overlapping into the active part of *S* are rendered (green).

### Simulation

The core of our simulation is an artificial space *S*^*N,N*^, with dimension *N* = *s*_px_ * *s*_im_, where *s*_px_ denotes the pixel size and *s*_im_ denotes the image size in pixels. *S* is thus a sub-lattice of the image *I* with nanometer resolution. It represents one frame with only a small amount of emitters in the fluorescent ON state. These emitters are simulated with the following properties: A spatial position in x- and y-direction *L*_*x*_, *L*_*y*_ ∈ [0, *N*], a lifetime distribution (Poisson, *t* = 90), a switch-on countdown (Poisson, *t* = 90) and a photon count (ph = randint(800,1500), Gaussian distribution *σ* = 0.2 ph). We define an empty subset of *L*_ON_ describing points in a fluorescent ON state. *L*_ON_ is recalculated each line, adding emitters based on a Poisson distribution 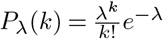 and deleting those which returned to a dark state. We further divide *S* into horizontal lines of size *s*_px_, representing the rasterization process of the detector (blue line in Figure 1 b). A time variable *t* is increased with each horizontal line by Δ*t* = 6 ms. The timestep of column-wise movements is neglected in our simulations since it is one order of magnitude faster.

If the switch ON countdown of a localisation is larger than zero, it is decreased by Δ*t* and the localisation is not rendered in the current line. If the variable drops below zero, the localisation is rendered and Δ*t* is subtracted from the remaining lifetime. Localisations surpassing their lifetime are not rendered in the subsequent lines and are deleted from *L*_ON_. The rendering process adds a crop of the PSF, corresponding to the localisations relative x- and y-position, to *S*. This crop equals the line width in y-dimension and is multiplied by the photon count of the emitter. A localisation has to be rendered for at least 40% of its ON-time to be accepted as a true positive. For the PSF shape, we use an airy disc model from astropy (14) with a varying radius *r* ∈ [525, 555]. However, this kernel can be easily replaced by a measured PSF for further applications. The simulated emitters are incomplete at the top or bottom. This depicts the switching process into the fluorescent ON/OFF state during the acquisition and is a typical feature of FLIMbee measurements. Subsequently, the image is resized by opencv’s (15) InterArea interpolation to *s*_*im*_. Noise is added corresponding to (16). For our training we simulated 9 × 9 × 3 crops of SMLM data. Each crop contains *n* ∈ [0, 10] localisations.

### Reconstruction

Our reconstruction pipeline is composed of several steps. First, regions of interest are detected by a trainable wavelet based peak detection layer and cropped to a 9 × 9 × 3 patch around the detected maximum, taking the temporal context of the previous and subsequent frame into account. The selected crops are further processed in one of the network architectures described in the following sections, ultimately creating a feature space describing the predicted emitters. This feature space equals the original crop data in size and contains a stack estimating the positions Δ*x* and Δ*y* relative to the pixel center, the emitter intensity *N*, the corresponding uncertainties *σ*_*x*_, *σ*_*y*_, *σ*_*N*_, the probability *p* for a pixel to contain an emitter, and an estimation *B* of the local background. This output format as well as the loss function were adapted from DECODE (7). If an emitter is close to the edge of a pixel, i.e. Δ*x/*Δ*y* are close to 0.5, the corresponding probability is often distributed over two adjacent pixels. Therefore, we defined the following conditions to retrieve a localisation: If a classifier pixel value exceeds the given threshold, a cross shaped filter is applied. If the convolved pixel exceeds 0.7, the pixel with the highest value of that formation is accepted as localisation. If it exceeds 1.4, the pixel with the second highest value is also accepted.

### Trainable wavelet layer

To reduce the dimensionality of the reconstruction problem, we developed a binning that crops localizations to a ROI of 9 × 9 pixels. These ROIs are identified with a trainable wavelet filter bank. To ensure perfect reconstruction, deconstruction and reconstruction filters share the same weights. Orthogonality of the filter bank is provided by coupling the learning process of low pass *lp* and high pass *hp* filter banks with a Gram-Schmidt process:

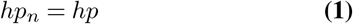

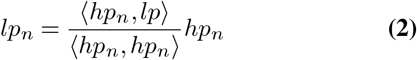

A bias followed by a ReLU activation is applied to the decomposed frequency images. A filtered image can then be reconstructed using the decomposed frequency images and the inverted filter bank. Given the noise and disruption-free ground truth as training data, the algorithm is able to filter spatial frequency components that are “PSF-like”. Potential localisations can then be identified by a local maximum detection. The denoised data can be of additional use for difficult reconstructions, e.g., when PSFs are shifted or disrupted as in FLIMbee measurements.

### Trainable CS layer

A major challenge of CS algorithms is the choice of suitable hyperparameters. We implemented the fast iterative shrinkage-thresholding algorithm (FISTA) (17), where the thresholding parameter λ significantly affects the number of iterations needed for convergence. Higher λ implies more background information to be filtered and leads to a faster convergence but can on the other hand lead to undetected localisations. Smaller λ implies less background and can lead to the detection of false positives. In classical approaches, λ is set globally dependening on the noise level. Since our method works with ROIs, an appropriate λ depending on the local noise level can be estimated. We implemented a classical CNN (Convolutional Neural Network) consisting of three convolutional layers followed by three dense layers, predicting a specific λ parameter for each crop (supplementary Fig. 7). Constraining this part of the network is challenging, as the network easily loses its gradient either by converging to a high λ, resulting in a zero output, or by converging to zero, resulting in no benefit of the compressed sensing operation. Therefore, it is crucial to regularize this step and to use suitable layer initialization. The λ estimation part of our network is followed by a sigmoid activation, multiplied by 0.025 corresponding to the maximal λ for a noiseless image that does not result in zero as output. Dense layers are initialized with a random normal distribution of mean *μ* = 0.5, standard deviation *σ* = 0.3 and truncated normal bias. We further implemented a functional test displaying the output of the first inception layer together with the estimated λ and the CS-reconstructed sub-lattice image to monitor the λ estimation process.

### Network architecture

To combine CS and artificial intelligence we implemented and evaluated the following network architectures:

#### CS CNN

Our first approach for a network design was to use a simple CNN as shown in Figure 2 a. We used the aforementioned FISTA layer as a prior, downsampled the sub-lattice back to the input dimensions, concatenated the original input image, and applied a set of convolutions to generate the eight described feature space maps.

**Fig. 2.**
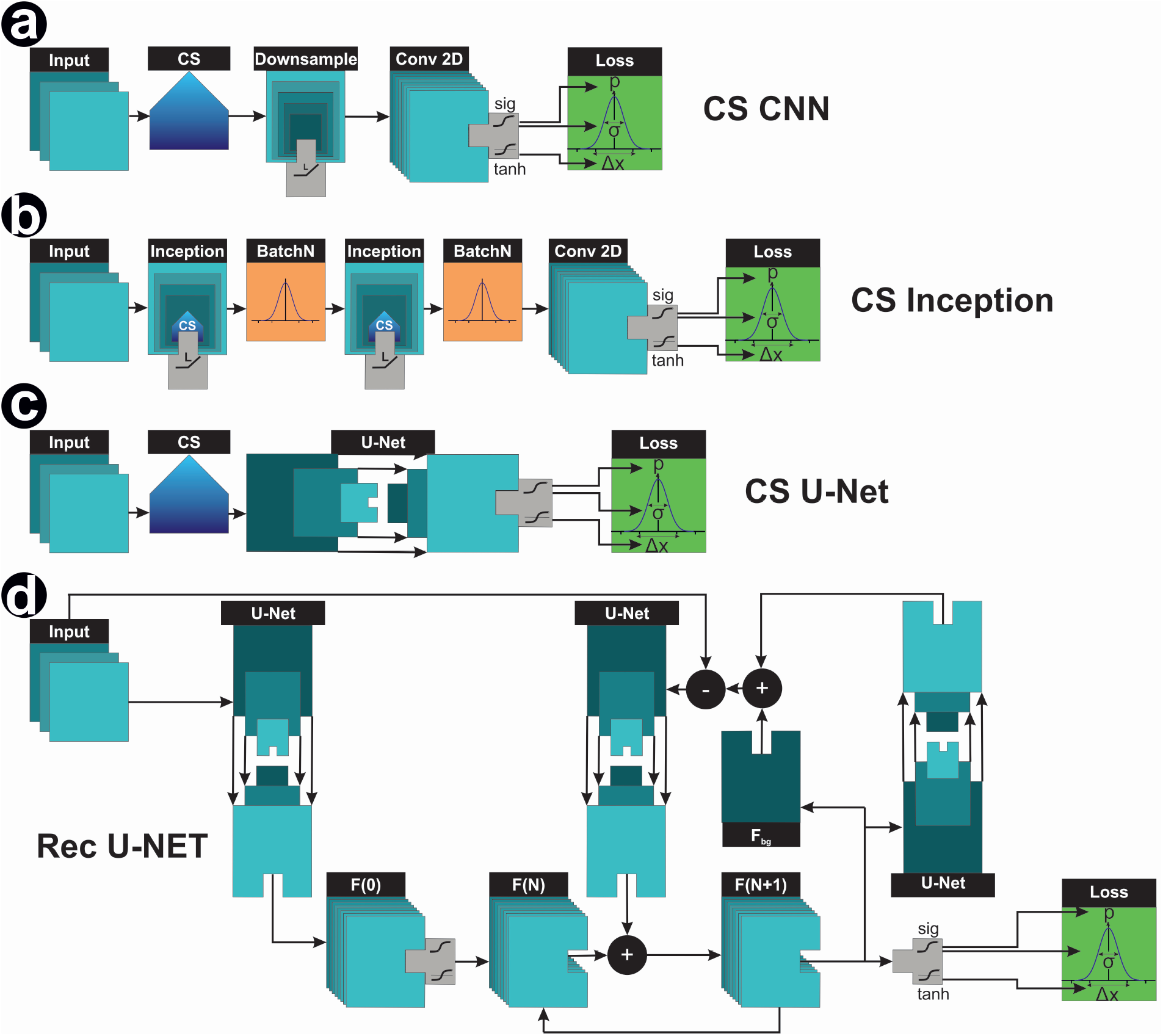
Network models. Image data containing the temporal context of the previous and subsequent frame are processed in different network models. (a) CS CNN uses compressed sensing as a prior and applies several convolutional layers. (b) CS Inception integrates the CS component deeper into the neural network. (c) CS U-Net uses compressed sensing as a prior and computes the feature space with a U-Net architecture (d) Rec U-Net aims to unroll the CS algorithm with iterative encoding and decoding from image to feature space and vice versa. For all network models the feature space is processed with sigmoid and tanh activations and fed into a Gaussian mixture model to compute the loss.

#### CS Inception

In a more sophisticated approach (Fig. 2 b), we integrated the concept of CS deeper into machine learning rather than using it only as a prior for computation. We built an architecture (Fig. 7) similar to Inception (18). The aim is to run a first inception layer with a very low CS iteration count as a prior for a second inception layer with a higher iteration count. While inception layer 1 can focus on improving the image quality, inception layer 2 can reconstruct coordinates with a lower error rate, i.e. compute a higher λ, resulting in faster convergence. The output of inception layer 2 is processed in a convolutional path similar to the first approach, reconstructing to eight feature layers in the original image dimensions.

#### CS U-Net

The U-Net (19) is a widely used neural network architecture for image-to-image tasks and was for example used in DECODE (7). The dimension of the input is step by step reduced in a down-sampling path, ultimately resulting in a dense feature space. In the subsequent up-sampling path, the dense information is combined with the corresponding layers of the down-sampling path containing spatial information(Fig. 2 c).

#### Recursive U-Net

A recent promising approach replaced the high iteration count of classical compressed sensing with ReLU activated convolutional layers (20). The feature space is connected to the sub-lattice via downsampling layers. The feature space is updated with additional details in each iteration. Therefore, the current estimation *x*(*t* + 1) is generated as *BN* (*x*(*t*) + *x*(0) + *update*), where *BN* describes a batch normalization, similar to ResNet (21). Since it is difficult to constrain an output in a convolutional network as sparse and our aim is to reconstruct a feature space in the original image size, predicting the coordinate center pixel and offset, we propose an iterative encoding and decoding between feature and image space. The model can be seen in Figure 2 d. In an initial step, we compute a first estimation for the feature space *F* (0). This estimation is updated *F* (*N* + 1) = *F* (*N*) + *F*_update_ by encoding the feature space to image space, adding the noise estimation *F*_bg_, calculating the element wise difference to the original image and subsequently encoding the obtained deviation back to feature space for element-wise addition with the previous estimation.

### Activation

A detailed visualization of the activations is shown in Figure 6. We used a sigmoid function on the output slice of the classifier image, to map each output pixel to a probability *p* ∈ [0, 1]. We constrained *σ*_*x*_, *σ*_*y*_ and *σ*_*N*_ to ∈ [0, 3] with a sigmoid activation multiplied by three to limit the standard deviation to a reasonable interval. A tangens hyperbolicus activation was applied to the subpixel coordinates Δ*x* and Δ*y* to clip these values into the range of [−1, 1]. This is important to maintain the advantages of local reconstruction while neglecting localisations beyond the local context. We also considered softmax as activation function for the classifier image. Despite a higher learning rate, this approach has major drawbacks. Outputs always cover the full range *p* ∈ [0, 1]. This gives rise to false positives if there is no localisation within the observed region, or false negatives if more than one active localisation is present.

### Loss function

The loss function of our network is composed of several components and was adapted from DECODE (7). We implemented a localisation loss for predicted emitters, a count loss for an accurate number of localisations, a prediction probability close to one or zero, and a background loss predicting the noise level. The localisation loss represents a Gaussian mixture model of the probability *p*_*i*_ for every pixel to contain a localisation, the position of said localisation *x*_*i*_ = *x*_px_ + Δ*x, y*_*i*_ = *y*_px_ + Δ*y*, where *y*_px_, *x*_px_ denote the coordinates of the current pixel and Δ*x*, Δ*y* its value, the estimated intensity *N* as well as the estimated error *σ* for each variable:

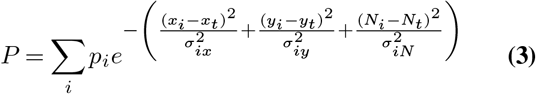

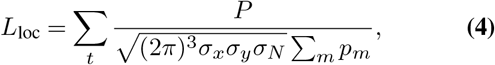

where *x*_*t*_ and *y*_*t*_ denote the ground truth coordinates and *N*_*t*_ the ground truth intensity. The count loss is

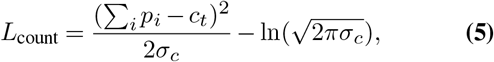

where *σ*_*c*_ = Σ_*i*_*p*_*i*_(1 − *p*_*i*_) encourages results close to 0 and 1 and therefore, reduces uncertainty. The background loss is

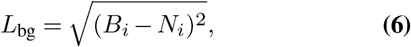

where *B*_*i*_ denotes the predicted background and *N*_*i*_ the noiseless ground truth. The total loss is the sum of the individual loss functions:

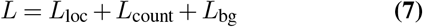

### Model Evaluation

To evaluate the performance of our approach, we use the RMSE (Root Mean Squared Error) and the Jaccard-Index *JI*:

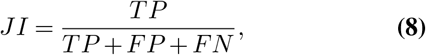

where *TP* are the true positives, *FP* the false positives and *FN* the false negatives. For real datasets with unknown ground truth, we used the Fourier Ring Correlation (FRC) as proposed in (22) (12)

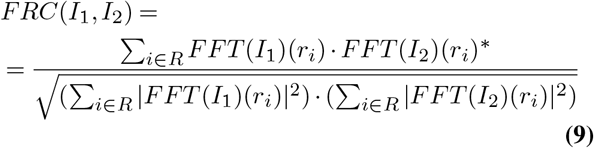

### Training Procedure

Using our simulation, we created a dataset consisting of 40 batches each containing 4 × 1000 crops. Three of these sub-batches are used for training and one for evaluation. While the noise simulations are completely random for each crop, sigma is different for each batch and in the range of *σ* ∈ [175, 185]. The network is trained for 150 iterations followed by one evaluation circle, where we compute the JI, the RMSE and a validation loss. We used an Adam optimizer with a learning rate of 10^−4^. Neural networks were implemented in Tensorflow 2 and trained on a Nvidia GTX 1080 TI GPU.

## Results

### Ground truth from simulated FLIMbee experiments

The first step before training a deep neural network is to obtain suitable ground truth. Training data can either be obtained from experiments and labeled by hand or by existing algorithms, or it can be generated using simulations. Since *d*STORM FLIMbee experimental data are difficult to measure and the performance of classical reconstruction algorithms is limited, we used the latter and developed a computer simulation of the FLIMbee measurement process. Using this tool, we simulated a dataset consisting of 40 batches, each containing 4 × 1000 crops. Three of these sub-batches were used for training and one for evaluation.

### Trainable wavelet filter to find regions of interest

Reconstructing SMLM data requires a lot of computational power, as each super-resolved image is composed of millions of localisations retrieved from thousands of frames. On top of that, compressed sensing algorithms are computationally expensive: The size of the reconstruction matrix and therefore the speed of the reconstruction scale with the fourth power of image size, *m*^4^. We reduce the dimension of the reconstruction problem by implementing a differentiable wavelet filter bank, trained to search for frequencies that resemble a PSF. ROIs are then cropped in a 9 × 9 × 3 area around the detected maximum, taking the temporal context of the previous and subsequent frame into account (Fig. 8).

### Deep neural networks with CS

To combine the advantages of CS with the benefits of artificial intelligence, we implemented and trained four different network architectures. The first approach was a simple CNN using the CS layer as a prior (Fig. 2 a). In the second approach, we used an Inception-like architecture (18) combined with CS. This network has two steps with different iteration counts and values of λ (Fig. 2 b). The third approach combines a classical U-Net architecture with CS, and the fourth network is a recursive U-Net-like architecture mimicking the iterative structure of CS. In all cases, the first layer has a size of 9 × 9 × 3 pixels, using the ROIs identified by the trainable wavelet transform as input.

To further constrain the compressed sensing part of our network we tried to implement an additional loss term for the CS layer to encourage a sparse reconstruction. For this term, we track the normalized compressed sensing output *b* of the inception layers and penalize entries differing from zero using the L_1_ loss. To prevent the compressed sensing part form diverging to zero, we apply the convolution matrix *A* to the CS output: *s* = *Ab*. In case of an optimal reconstruction, this operation convolves a sparse sub-lattice with the measurement function while downsampling to the original image size. The result *s* is a denoised version of the original image. The loss can then be computed as squared difference to the noiseless training data:

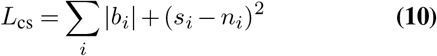

Since this did not improve the results significantly we discarded the loss term in our final network versions.

### Comparison of network architectures

We compared the performance of the four different network architectures on a separate simulated validation dataset. We computed the RMSE, JI and validation loss every 150 steps on an independent test batch during training (Fig. 3). The recursive U-Net achieved the best results compared to the other methods. It can be observed that CS in the form of FISTA is a solid prior for the Inception-like network, leading to an early increase in metrics. For higher iterations, however, the network metrics converge. The training time per epoch for the Inception architecture (2091.3 s) is 32.6 times higher than the training time per epoch of the CS-Res-U Net (64.1 s). For the evaluation of an example dataset, the Rec U-Net still introduces a 5.6 fold acceleration in comparison to the Inception architecture. A detailed evaluation of the training and evaluation performances is shown in Table 1.

**Table 1.** Training and inference time of different network architectures on a Nvidia GTX 1080 TI. Training times are measured per epoch. For the evaluation of inference time, a FLIMbee dataset with 4500 frames of 45×45 px is used.

| Network architecture | Training time [s] | Inference time [s] |
| --- | --- | --- |
| CS-CNN | 1102.0 | 440.8 |
| CS-Inception | 2091.3 | 502.5 |
| CS-U | 596.4 | 72.5 |
| CS-Res-U | 64.1 | 88.8 |

**Fig. 3.**
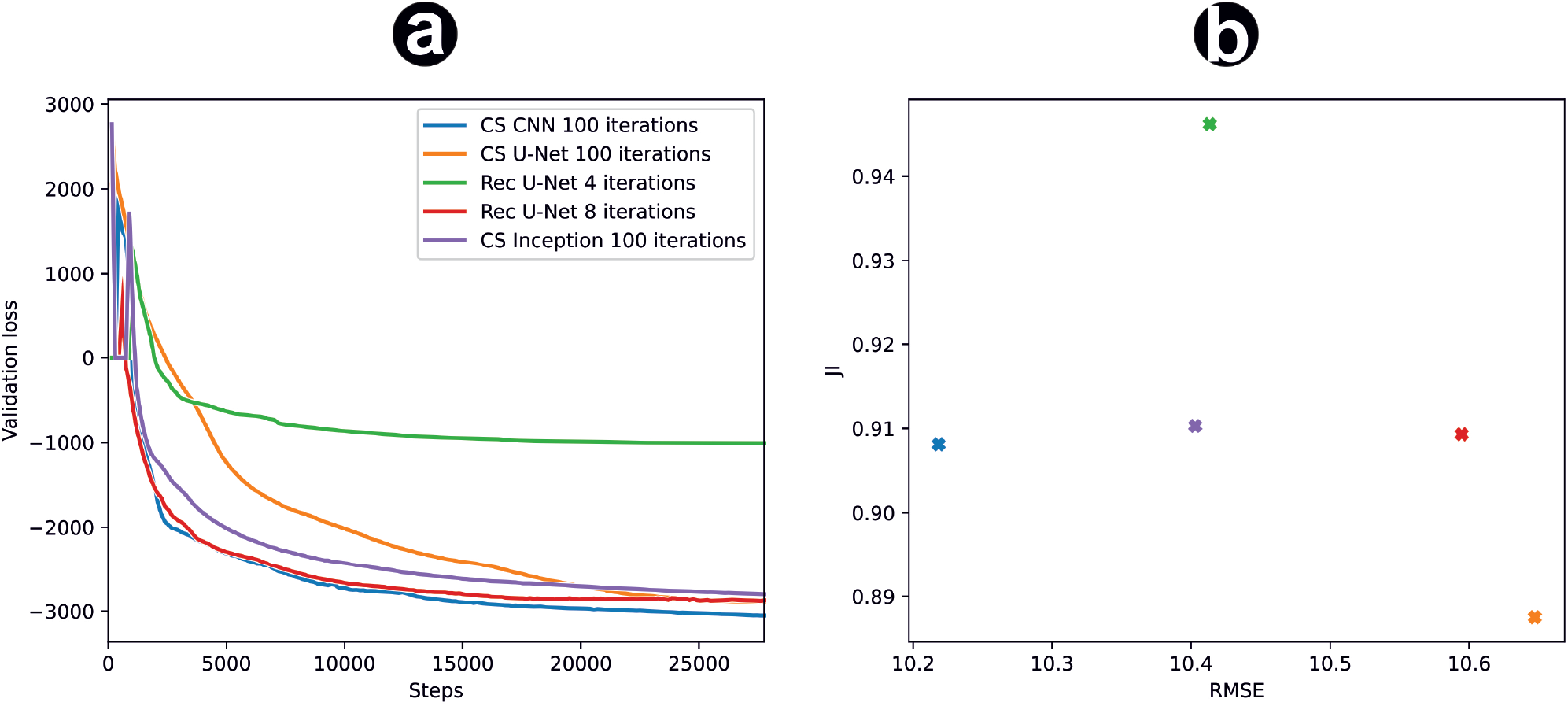
Comparison of different network architectures. (a) Validation loss over training steps. (b) Jaccard Index and RMSE of the tested models.

### Application to experimental data

We performed experiments with a FLIMbee galvanometric scanner as described in the methods (Fig. 4) and tested the trained networks on these real data. The raw data shows the typical interrupted PSF as well as a varying intensity between lines. Trained for these non-linearities, our network is able to precisely predict the center of these localisations. To assess the quality of prediction on experimental data, we used FRC and LineProfiler, as objective ground truth was not available in this case. The results (CSInception: 0.211 Rec U-Net: 0.265) indicate an improved reconstruction quality compared to classical fitters like ThunderSTORM (0.167). Note that an additional drift correction improves the quality of the reconstruction significantly. For the image in Fig. 4 we applied a linear drift. For the other images in Fig. 5, we used the ThunderSTORM RCC drift correction.

**Fig. 4.**
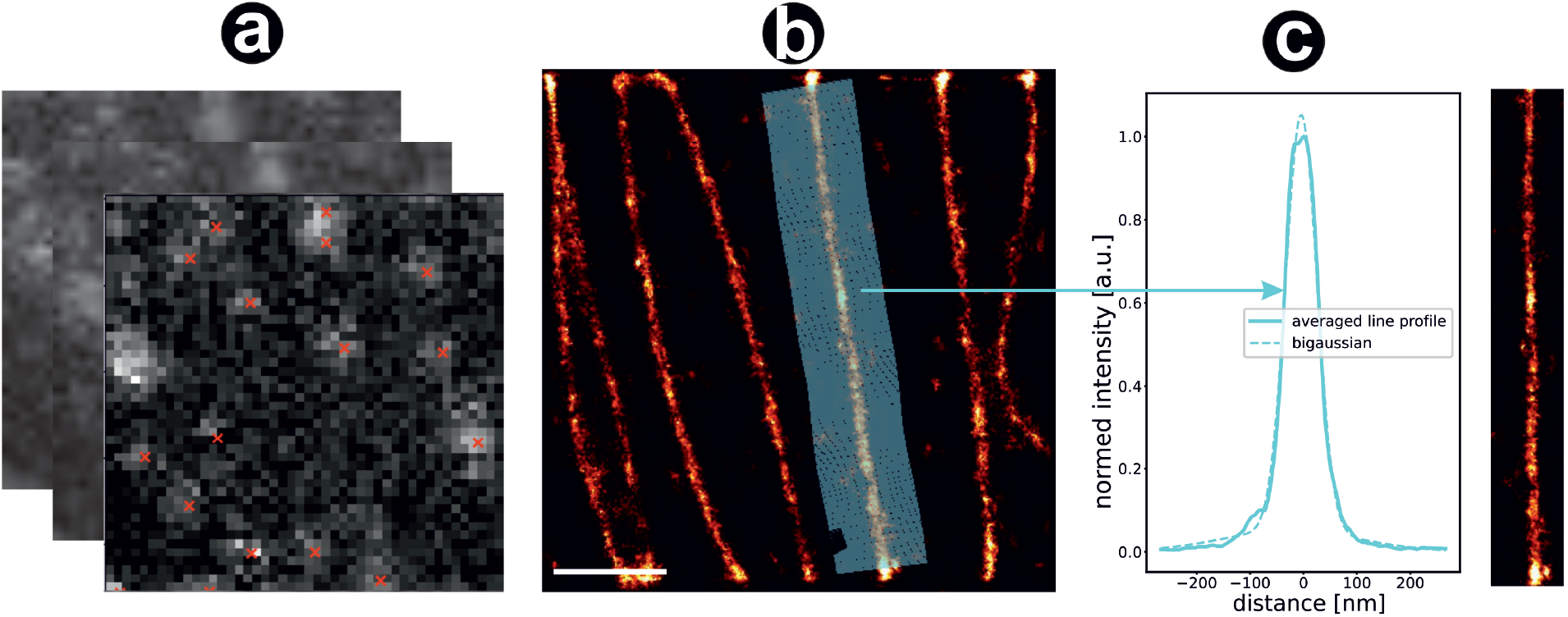
Reconstruction of FLIMbee *d* STORM microtubuli. (a) Fitting of disrupted PSFs with artificial intelligence. Red crosses denote the estimated location of a fluorophore. (b) Reconstructed super-resolution image. Scale bar = 10 μm (c) Line profile of the microtubule marked blue in (b).

**Fig. 5.**
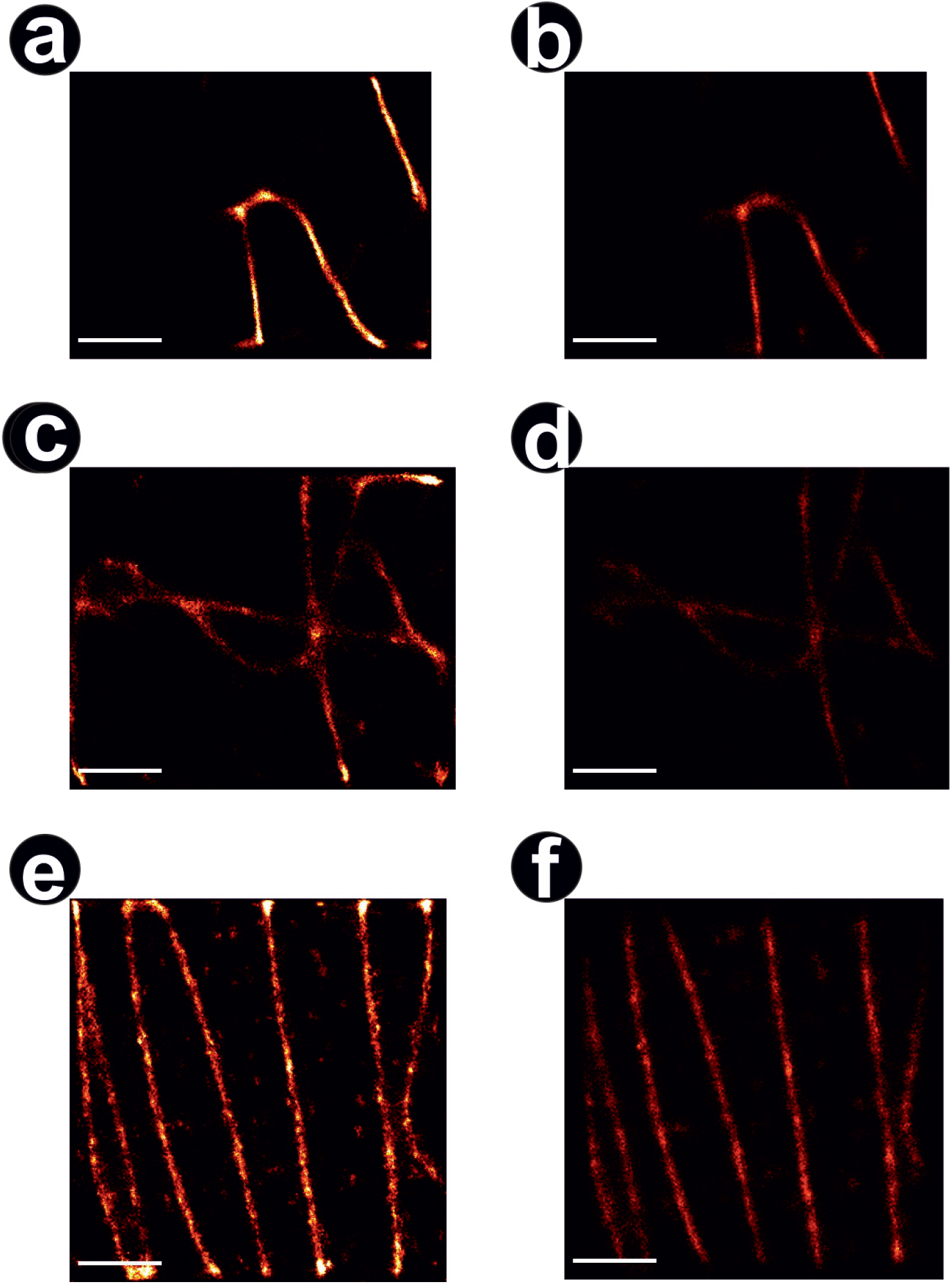
Comparison of FLIM data evaluated with thunderstorm and our method. Reconstructed images of AI (left) and Thunderstorm (right). Comparing the first and the second half of the localisation data, we obtain a Fourier Ring Correlation Coefficient (12) of 0.310; 0.187; 0.265 (left) and 0.179; 0.187; 0.167 (right) respectively. Scale bar = 10 μm

**Fig. 6.**
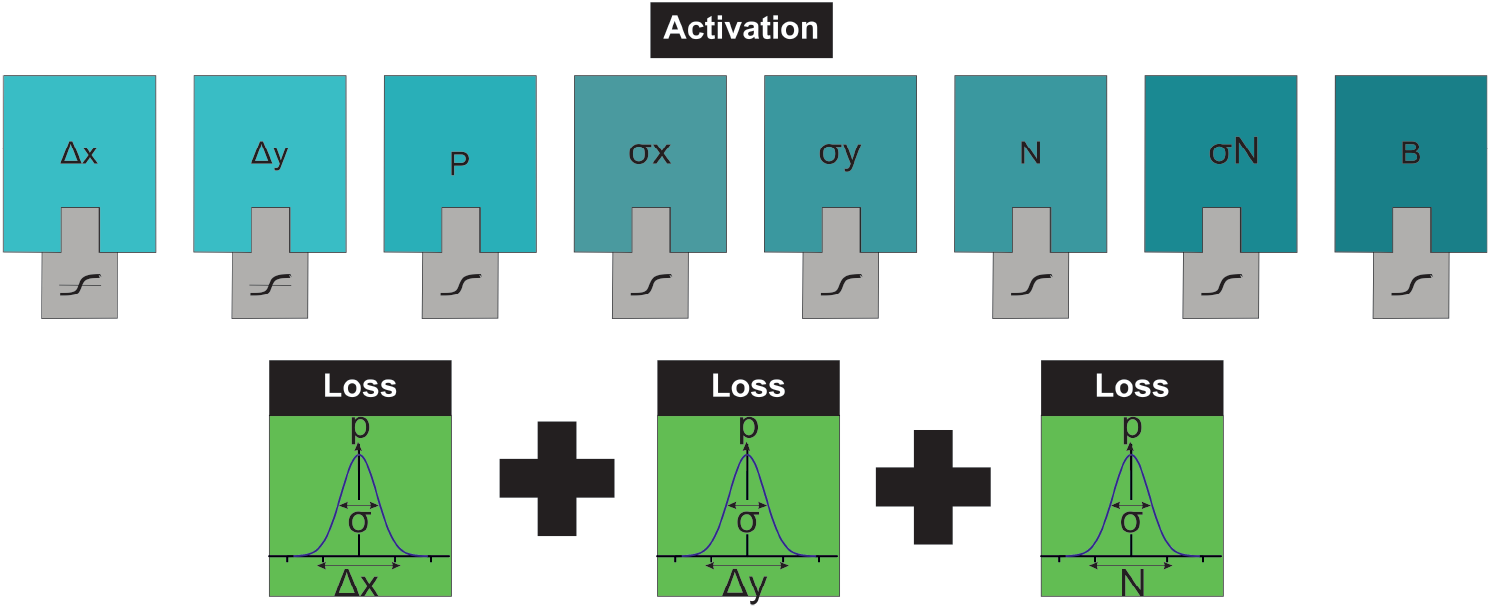
Activations of the output layer. The relative positions Δ_*x*_ and Δ_*y*_ are fed into a Gaussian mixture model with the local probability *p* and the positional uncertainties *σ*_*x*_ and *σ*_*y*_. The estimated intensity *N* is included into the localisation loss. While Δ_*x*_ and Δ_*y*_ are *tanh* activated, all other components are sigmoid activated.

**Fig. 7.**
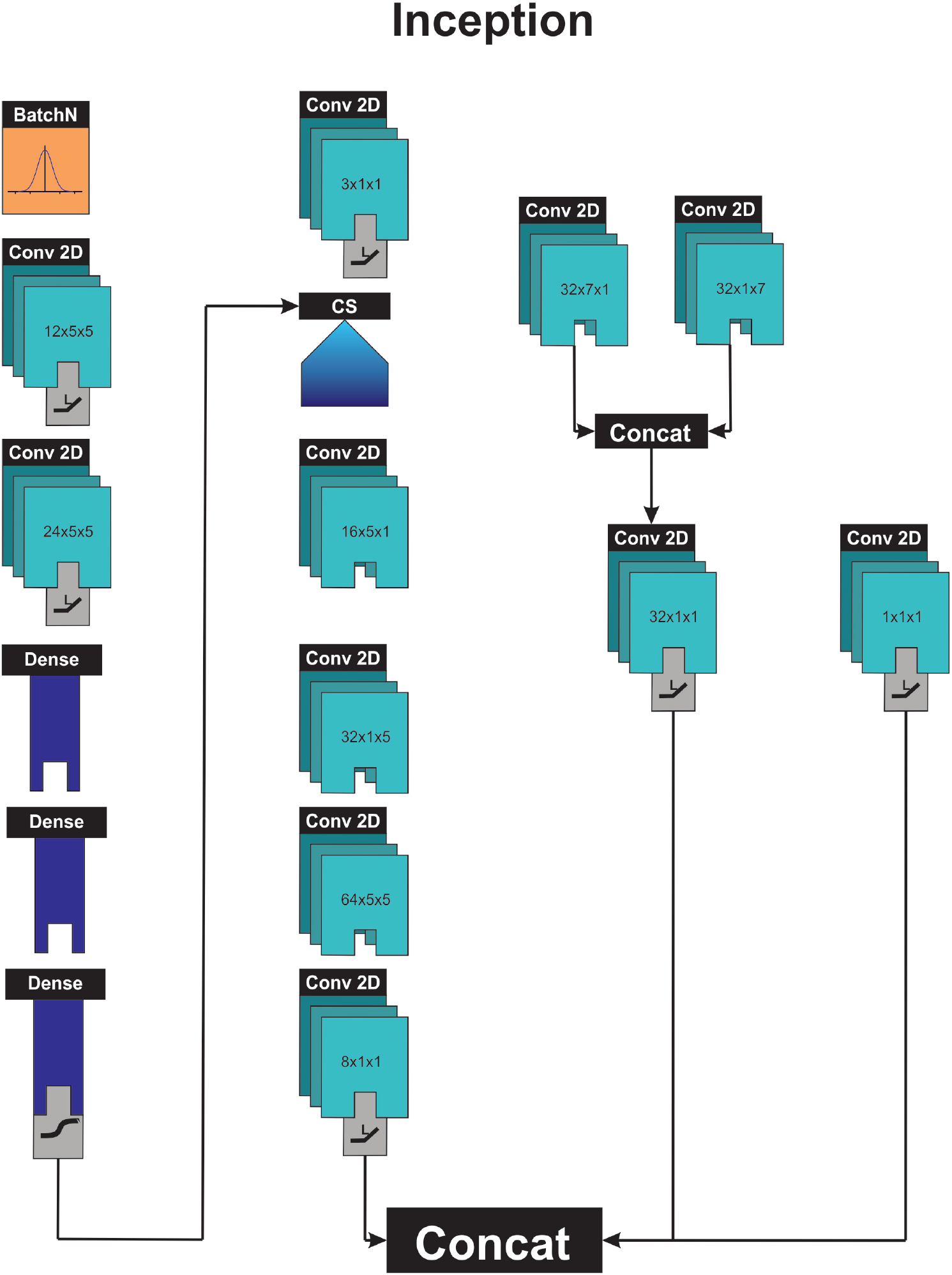
Inception building block. This building block is derived from the inception network (18). The input is processed in 4 different paths (from left to right). We estimate the compressed sensing parameter λ with a conventional CNN. The input is processed by a bottleneck layer, followed by the compressed sensing layer. Several convolutional layers restore the original image size. A feature detector applies asymmetric filters from different directions. A pass-through only applies an activation function, passing forward the original image.

**Fig. 8.**
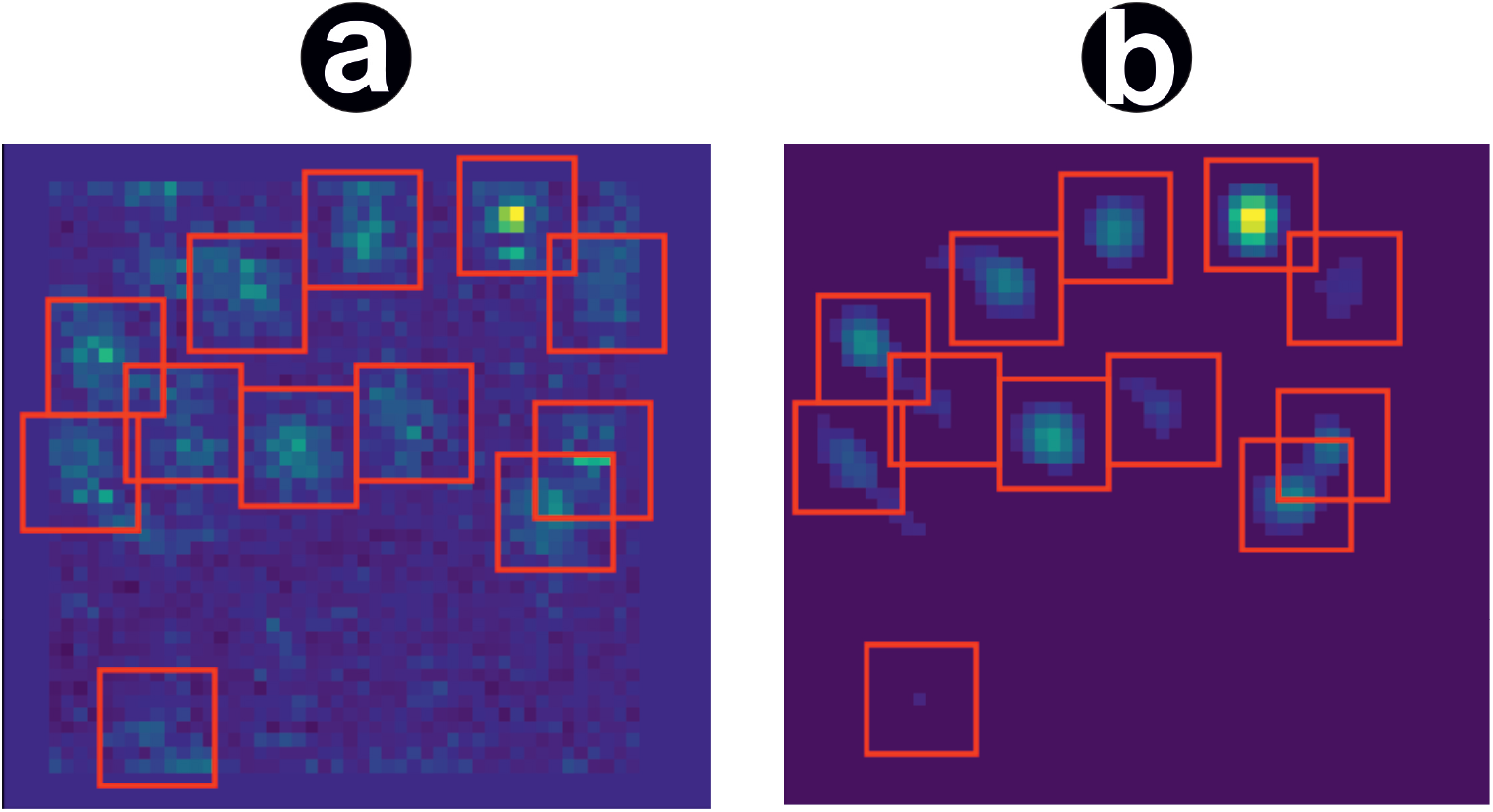
Wavelet filter bank peak detection on an example frame of a FLIMbee measurement. The input image (a) is deconstructed with a wavelet filter bank, trained to extract frequencies resembling a PSF. A threshold is applied before reconstructing to generate a denoised image (b). Potential emitters are identified with a local maximum detection. The detected peaks are marked with red rectangles.

## Discussion

We developed a robust data simulator for FLIMbee SMLM measurements, combining the method-specific disrupted PSFs with accurate noise simulations. We furthermore introduce a learnable wavelet filter that can be trained to accurately detect emitters and crop them to enhance the speed of the evaluation pipeline. Finally, we implemented and evaluated different approaches to integrate compressed sensing operations into deep neural networks for the reconstruction of super-resolution images from nonlinear disrupted PSFs. The Rec U-Net architecture achieved the best JI and RMSE performance on simulated data as well as the best FRC score on real data, while architectures with CS-like sparse representations did not perform as well. This indicates that sparse representations might not be optimal for the learning process of neural networks. A possible reason may be the large amount of zero values in the sparse representation, leading to a vanishing gradient for large fractions of the feature space. Neural networks might be better suited to create a parameterized representation of the sparse sub-domain, like compressed sparse row (23), or to directly compute the feature space representation as proposed in the Rec U-Net.

The trainable wavelet filter is an efficient way to identify regions of interest in SMLM data. Trained on realistic simulated data, it is able to filter background frequencies and to accurately determine regions of interest. Fitting the center of sparsely activated emitters is a redundant problem, so preselecting regions of interest has several advantages over reconstructing a whole image. The subsequent reconstruction network is scalable, since the reconstructed regions of interest always have identical size. On top of that, training duration and network depth can be reduced drastically. However, there are cases where prefiltering has its limits or even introduces disadvantages. High density samples pose a problem since emitters overlap and have less resemblance with the original PSF. Data that diverges too much from the original training data can also lead to loss of localisations. For the given problem of low density emitters with disrupted PSFs, however, it is an efficient way to identify regions of interest.

As stated in (7), inaccurately estimated localisations tend to be reconstructed towards the center of a feature space pixel. This can be overcome by adjusting the precision threshold and/or the reconstruction method (local maxima). Interestingly, this feature was also observed for classical compressed sensing methods like (24) and seems to be a general problem of discrete feature spaces.

The fact that metrics converge at high iterations may be caused by the linear convergence of L1 minimization algorithms, as the available information before full convergence is limited. Another possible explanation is that the training process is able to extract the necessary information even with low iteration counts. Interestingly, the best results in terms of JI and RMSE do not coincide with the best validation loss. Possible explanations of this behavior include overfitting, or local minima with very low underestimated localisation uncertainty.

As can bee seen from Table 1, the initial FISTA layer introduces a large computational cost, making the Rec U-Net approach much faster than the CS-like implementations. This architecture resembles the unfolding of CS interations in a deep neural network, as proposed by Gregor and LeCun (25). An approach, that has already been applied to SOFI super-resolution imaging using sparsity in the correlation domain (26) by unrolling the iterative FISTA compressed sensing algorithm into a deep neural network. Our results confirm that algorithm unfolding is an efficient method to combine the advantages of iterative compressed sensing methods with deep learning for reconstruction of high-resolution microscopy data and will likely see many other applications in the future.

The current state-of-the-art for fitting classical localization data according to the SMLM challenge (10) is DECODE (7). In this work, three independent U-Nets were applied to three consecutive frames to detect localizations. It was not possible to perform a direct comparison of our approach to DECODE, since its training process is tightly coupled to a simulation of frames with a spline-parameterized PSF that is not compatible with our confocal *d*STORM data. If the training process could be adapted to incorporate our simulator, it would be interesting to see if it performs as well as our CS U-Net-based approach, since we adapted our loss function and output format from DECODE.

## Conclusions

We developed a data generator for nonlinear PSFs in the context of super-resolved confocal lifetime imaging and were able to reconstruct localisations with improved accuracy compared to classical fitters by developing and training an artificial neural network. Next to an improvement in computation time, we demonstrate the adaptation of compressed sensing to deep neural networks for reconstructing non-linearly varying PSFs. Our results indicate that using a deep architecture like inception is beneficial to the models performance. Including local context by reconstructing to the original crop size as well as including the temporal context of the previous and subsequent frame improves the reconstruction quality significantly. Implementing compressed sensing into artificial neural networks is a promising concept, but further work has to be done to improve the implementation details. For an optimal solution, the CS part should fully converge. This is, however, computationally demanding since every iteration contains a nontrivial derivation used by the network for back-propagation. In comparison, algorithm unfolding appears to be a more efficient way to integrate compressed sensing and deep learning.

## ACKNOWLEDGEMENTS

MS has received funding from the European Research Council (ERC) under the European Union’s Horizon 2020 research and innovation programme (grant agreement No 835102) and the Deutsche Forschungsgemeinschaft (DFG SA829/19-1). PK has received funding from Deutsche Forschungsgemeinschaft (DFG KO3715/5-1).

## Author’s contributions

PK, MS and SR designed the project. SR wrote the code and performed the evaluations. DAH and DB performed the FLIMBee measurements. PK, SR and DAH wrote the manuscript. All authors read and approved the final manuscript.

## Software

The described software is available at the following github repository: https://github.com/super-resolution/ReCSAI.

